# AutoSpill: A method for calculating spillover coefficients to compensate or unmix high-parameter flow cytometry data

**DOI:** 10.1101/2020.06.29.177196

**Authors:** Carlos P. Roca, Oliver T. Burton, Teresa Prezzemolo, Carly E. Whyte, Richard Halpert, Łukasz Kreft, James Collier, Alexander Botzki, Josef Spidlen, Stéphanie Humblet-Baron, Adrian Liston

## Abstract

Compensating in classical flow cytometry or unmixing in spectral systems is an unavoidable challenge in the data analysis of fluorescence-based flow cytometry. In both cases, spillover coefficients are estimated for each fluorophore using single-color controls. This approach has remained essentially unchanged since its inception, and is increasingly limited in its ability to deal with high-parameter flow cytometry. Here, we present AutoSpill, a novel approach for calculating spillover coefficients or spectral signatures of fluorophores. The approach combines automated gating of cells, calculation of an initial spillover matrix based on robust linear regression, and iterative refinement to reduce error. Moreover, autofluorescence can be compensated out, by processing it as an endogenous dye in an unstained control. AutoSpill uses single-color controls and is compatible with common flow cytometry software, but it differs in two key aspects from current methods: (1) it is much less demanding in the preparation of controls, as it does not require the presence of well-defined positive and negative populations, and (2) it does not require manual tuning of the spillover matrix, as the algorithm iteratively computes the tuning, producing an optimal compensation matrix. Another algorithm, AutoSpread, complements this approach, providing a robust estimate of the Spillover Spreading Matrix (SSM), while avoiding the need for well-defined positive and negative populations. Together, AutoSpill and AutoSpread provide a superior solution to the problem of fluorophore spillover, allowing simpler and more robust workflows in high-parameter flow cytometry.

## Introduction

Fluorescently-labeled antibodies and flow cytometry have been the workhorse for single-cell data generation in many fields of the biosciences since its development in the late ’60s^1^. The ability to rapidly collect quantitative data from millions of single cells has driven the understanding of heterogeneity in complex cellular mixtures, and led to the development of many fluorescence-based functional assays^2–5^. The diverse utility of flow cytometry has driven constant demand for an expansion in the number of parameters to be simultaneously measured. Development of novel fluorophores and advances in laser technology have provided a steady increase in the number of parameters that can be measured on state-of-the-art machines, roughly doubling each decade since the ‘70s (“Roederer’s Law for Flow Cytometry”)^6^.

The development from single-color flow cytometry to ultra high-parameter flow cytometry has allowed an enormous growth in the data collected per cell. In our own field of immunology, high-parameter flow cytometry panels have become necessary, with multiple markers required to identify cellular lineages, major subsets, and activation markers. A key limitation with high-parameter flow cytometry, however, is the spectral overlap of fluorescent dyes^7^. This results in the spillover of fluorescence to detectors different from the detector assigned to each dye (in classical flow cytometry). Removing this unwanted spillover, i.e. compensating, is a necessary preliminary step in the data analysis of multi-color flow cytometry.

State-of-the-art flow cytometers, with ~30 channels, make compensation increasingly difficult as the number of channels grows, due to the unavoidable overlap between emission spectra of fluorescent dyes. The difficulty of experimental design has followed the growth in fluorophore options, to the point where the development, refinement, and validation of ultra-high parameter panels can take months to years of expert input^4,8-10^. Indeed, the development of mass cytometry as an alternative technology is largely driven by its lack of spillover^11^, as otherwise the technology compares unfavorably to flow cytometry in several aspects^6^.

Unlike the extensive development efforts in fluorophore generation, fluidics refinement, and laser addition, the basis for dealing with spillover in flow cytometry has largely remained unchanged. Current compensation algorithms are based upon the algorithm for spillover calculation proposed by Bagwell and Adams, when flow cytometers worked with only a few fluorophores^12^. These approaches provide an estimation of the spillover matrix, in which the degree of spectral spillover between channels is estimated from single-color controls. A compensation matrix is obtained by inverting the spillover matrix, by which spillover is compensated out from experimental datasets. While effective in low-parameter datasets, where spillover is moderate to start with, in the case of high-parameter data this method often requires manual adjustment before proceeding with downstream analyses. This manual tuning entails manipulating a matrix with several hundred coefficients, which can be extremely challenging and time-consuming, thus severely constraining panel design in practice. In spectral flow cytometry, unmixing is carried out in a different way, but obtaining the spectral signature of each fluorophore is also based on single-color controls, and it still requires a similar calculation to estimate the spillover to every detector. In both cases, these approaches require single-color controls with well-defined positive and negative populations, which often forces the single-color controls to differ from those of the actual panel, increasing the complexity of the experiment.

We have developed a new algorithm, AutoSpill, to compensate flow cytometry data. This approach uses single-color controls, making it compatible with existing datasets and protocols. Unlike other compensation approaches, however, it calculates spillover coefficients by means of robust linear models. This method produces better estimation of spillover coefficients, without requiring well-defined positive and negative populations. Moreover, AutoSpill uses this improved estimation of the spillover matrix only as the initial value for an iterative algorithm that automatically refines the spillover matrix until achieving, for practical purposes, virtually perfect compensation for the given set of controls. In addition to providing optimal spillover matrices for compensating (or unmixing in spectral systems), and given that AutoSpill does not rely on well-defined positive and negative populations, it can calculate the autofluorescence spectrum of cells by treating it as an extra endogenous dye. Thus, it allows effective detection and removal of autofluorescence from experimental data.

A linear modeling approach can equally be used to estimate the increase in fluorescence noise or spread caused by compensating spillover. Thus, we also propose a second novel algorithm, AutoSpread, which calculates spillover spreading coefficients with linear models, thereby providing a Spillover Spreading Matrix (SSM) without the need for well-defined positive and negative populations in the single-color controls.

Together, AutoSpill and AutoSpread remove limiting constraints of traditional compensation methods, easing the preparation of compensation controls in high-parameter flow cytometry, making errors less likely, and facilitating the practical implementation of ultra high-parameter flow cytometry. AutoSpill is available through open source code and a freely-available web service (https://autospill.vib.be). AutoSpill and AutoSpread are available in FlowJo v.10.7.

## Results

### Tessellation allows robust gating

A critical first step in the processing of flow cytometry data is the elimination of cellular debris and other non-cellular contamination. This stage is typically performed by manual or automated gating of particles with the expected size and granularity, based on forward scatter and side scatter. In order to develop a fully automated pipeline, we sought to encode this initial cellular gating in the AutoSpill algorithm. After numerous tests on data provided by collaborating immunologists, we settled on a multi-step process with two tessellations, which demonstrated the required features of robust cell or bead identification. Figure 1 shows the initial gating for one single-color control of each set of controls. The multi-step process robustly identified the cellular fractions as desired, regardless of the presence of high amounts of cellular debris in the HS1 and HS2 datasets (Fig. 1, second and third column). It also worked correctly with beads (Be1 dataset), which exhibited substantially different forward-scatter/side-scatter profiles (Fig. 1, fourth column). For all channels and all datasets, the gate selected the cell/bead population in the desired density maximum, without needing manual adjustment.

**Figure 1:**
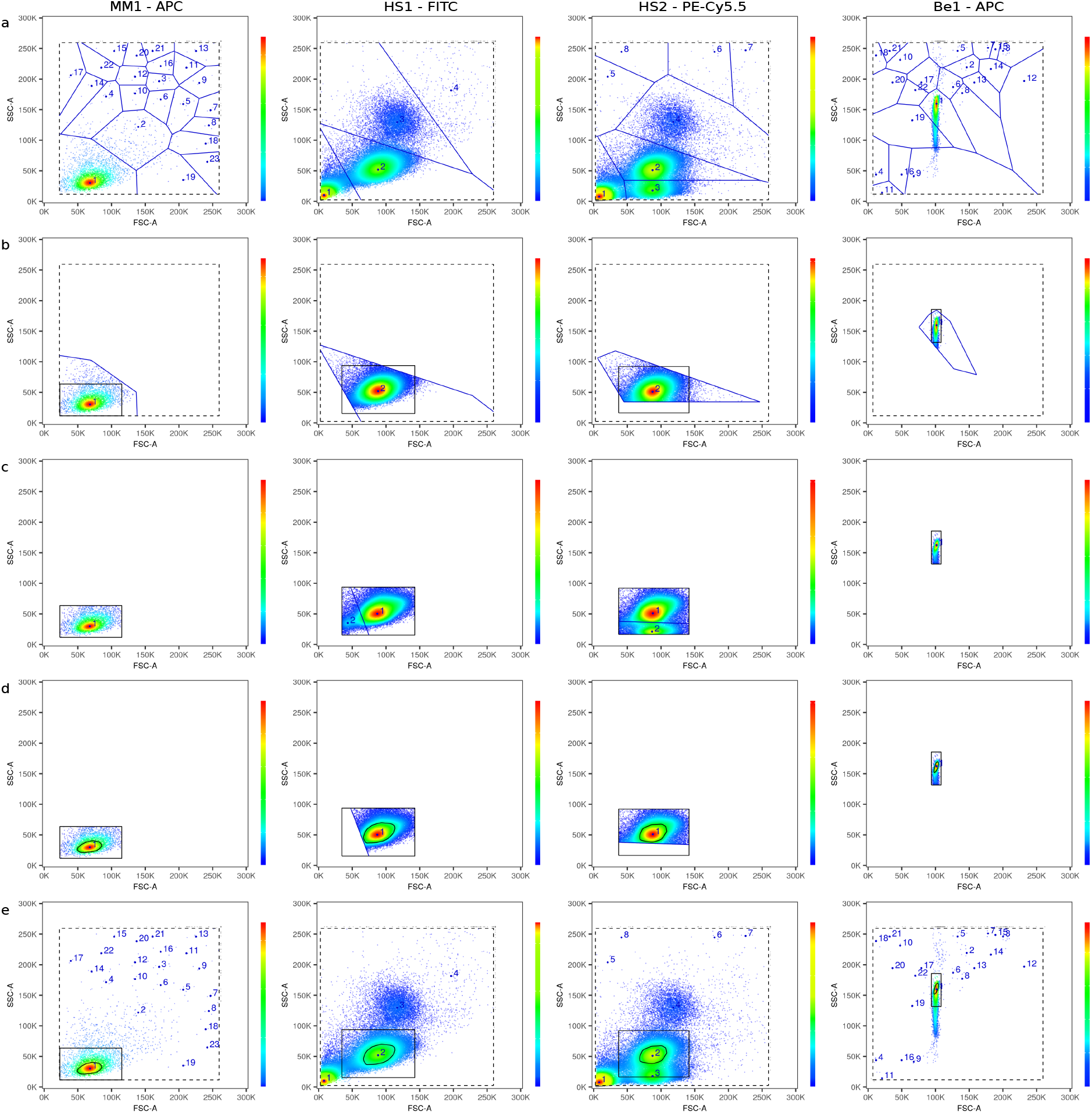
Tessellation allows robust initial gating. Results of gating before the calculation of compensation, using forward and side scatter parameters (as shown in the axes), for different samples with cells or beads. Columns show one gate example for each dataset, as indicated. Rows show the successive steps of the algorithm for each example: (a) bound calculation (dashed black line) and first tessellation (in blue), to identify the density maxima (blue points, with numbers showing decreasing order of density value); (b) region identification (solid black line) around the target maximum; (c) second tessellation (in blue), to isolate the target maximum from close maxima inside the region (point color and number as in (a)); (d) calculation of the boundary gate (black closed curve), by a threshold on density and a convex hull; (e) gate summary provided to package/website users, with same line, point, and color code as in (a)–(d).

### Robust linear regression effectively estimates spillover coefficients

The estimation of spillover coefficients is based on the comparison between the level of fluorescence detected in the primary channel (i.e. the detector dedicated to the dye or fluorophore, in classical systems, or the detector with highest signal, in spectral systems) and the secondary channels (i.e. every other detector). The linear relationship between the fluorescence levels of primary and secondary channels is not visible in the usual bi-exponential scale (Fig. 2, first and third column), but it becomes apparent in linear scale (Fig. 2, second and fourth column). The linear relationship between the primary and secondary channels shows that the ratio of fluorescence between the two channels is constant across a broad range of fluorescence levels. Thus, a linear regression can be used to properly identify the slope between the two channels, that is, the spillover coefficient. This approach produces a similar result to that achieved by the calculation of a slope between the median values of the positive and negative populations^12^, which is the method usually employed (Fig. 2, first and second column). Notably, however, the use of linear regression also allows the robust calculation of the slope in cases that the traditional approach was not designed to deal with: low numbers of positive events (Fig. 2b), without a well-defined positive population (Fig. 2c), or without well-defined positive and negative populations (Fig. 2d). The quality of compensation can be evaluated by the difference between the obtained compensation and the ideal one, with perfectly compensated data showing an exactly vertical distribution of data along the primary fluorophore (i.e. zero slope). While traditional estimation of spillover was successful to some extent in producing low-error compensation, in particular when distinct positive and negative populations were present (Fig. 2a, first and second column), errors were identified in particular channels, especially when populations did not conform to good separation (Fig. 2c, first and second column). In all cases, linear regression resulted in less compensation error (Fig. 2, third and fourth column).

**Figure 2:**
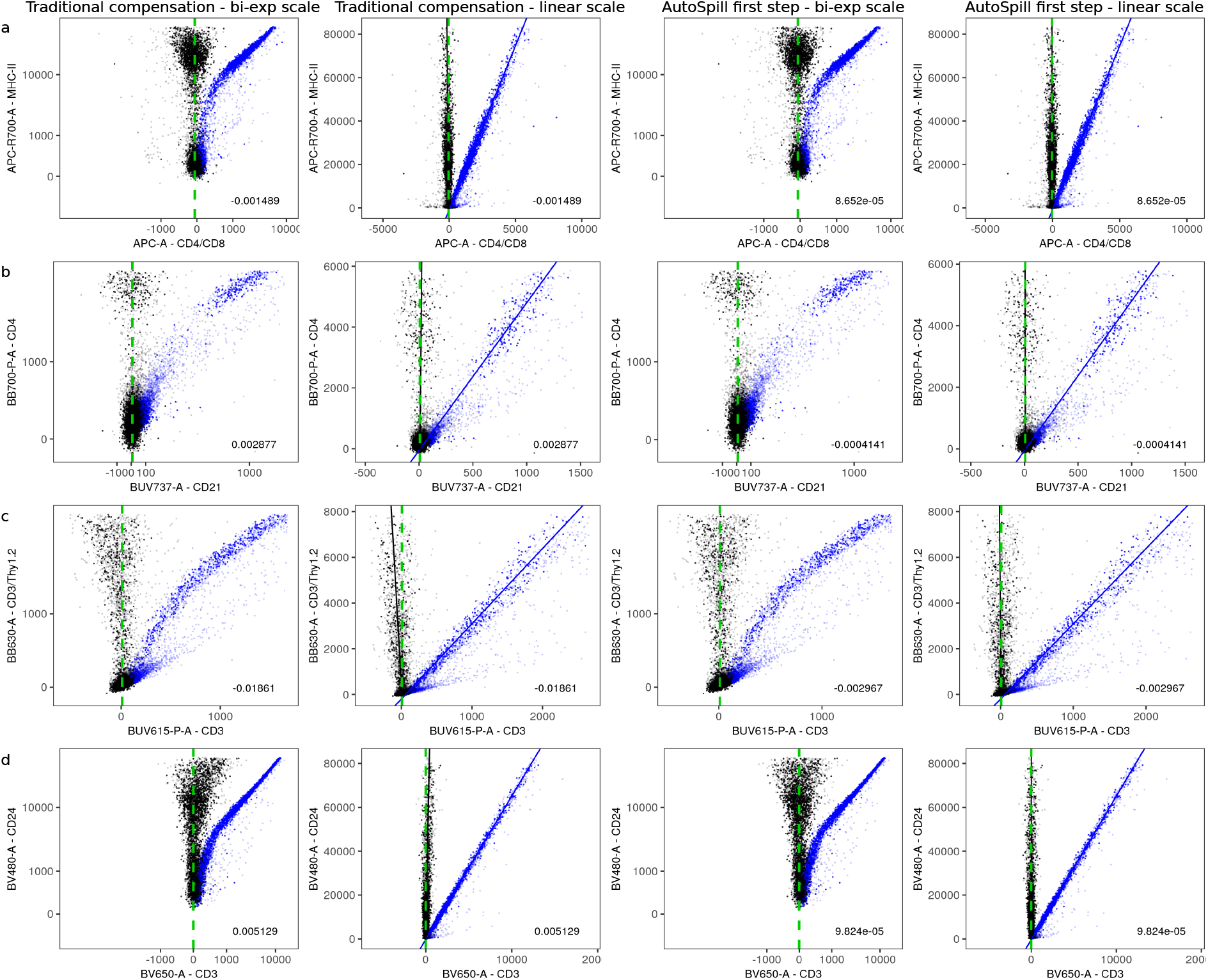
Robust linear regression effectively estimates spillover coefficients. Each row (a–d) shows a compensation example from the MM1 dataset, with the primary and secondary channels indicated, respectively, in the *y*-axes and *x*-axes. Compensation results are displayed using positive and negative populations (first column, bi-exponential scale; second column, linear scale), and robust linear regression (third column, bi-exponential scale; fourth column, linear scale). The linear relationship between the levels of fluorescence is not visible in bi-exponential scale, but it is very clear in linear scale. Uncompensated data is displayed in blue and compensated data in black. Dim points correspond to gated-out events, not used in the calculation. Lines in the second and fourth column (linear scale) show regressions of uncompensated (blue) and compensated (black) data. The slope coefficient of the latter provided the compensation error (number at the bottom right of each panel). Vertical green dashed lines are shown as a reference for perfectly compensated data.

### Iterative reduction of compensation error yields optimal spillover coefficients

The spillover coefficients obtained in the first iteration step by robust linear regression produced low-error estimates of the spillover matrix for all channels (Fig. 3). While this error level outperformed that of the traditional approach (Fig. 3), some channels exhibited a degree of overcompensation or undercompensation. While such errors are small, they nonetheless produce overcompensation or undercompensation noticeable in bi-exponential scale, which visually amplifies fluorescence levels close to zero. In a high-parameter flow cytometry panel, with multiple fluorophores present on small subpopulations, such errors can accumulate to the point of making individual channels effectively unusable. We therefore developed an iterative reduction of error, by successively obtaining better spillover matrices and compensated data. This iterative refinement of the spillover matrix reduced the compensation errors to negligible values (Fig. 3).

**Figure 3:**
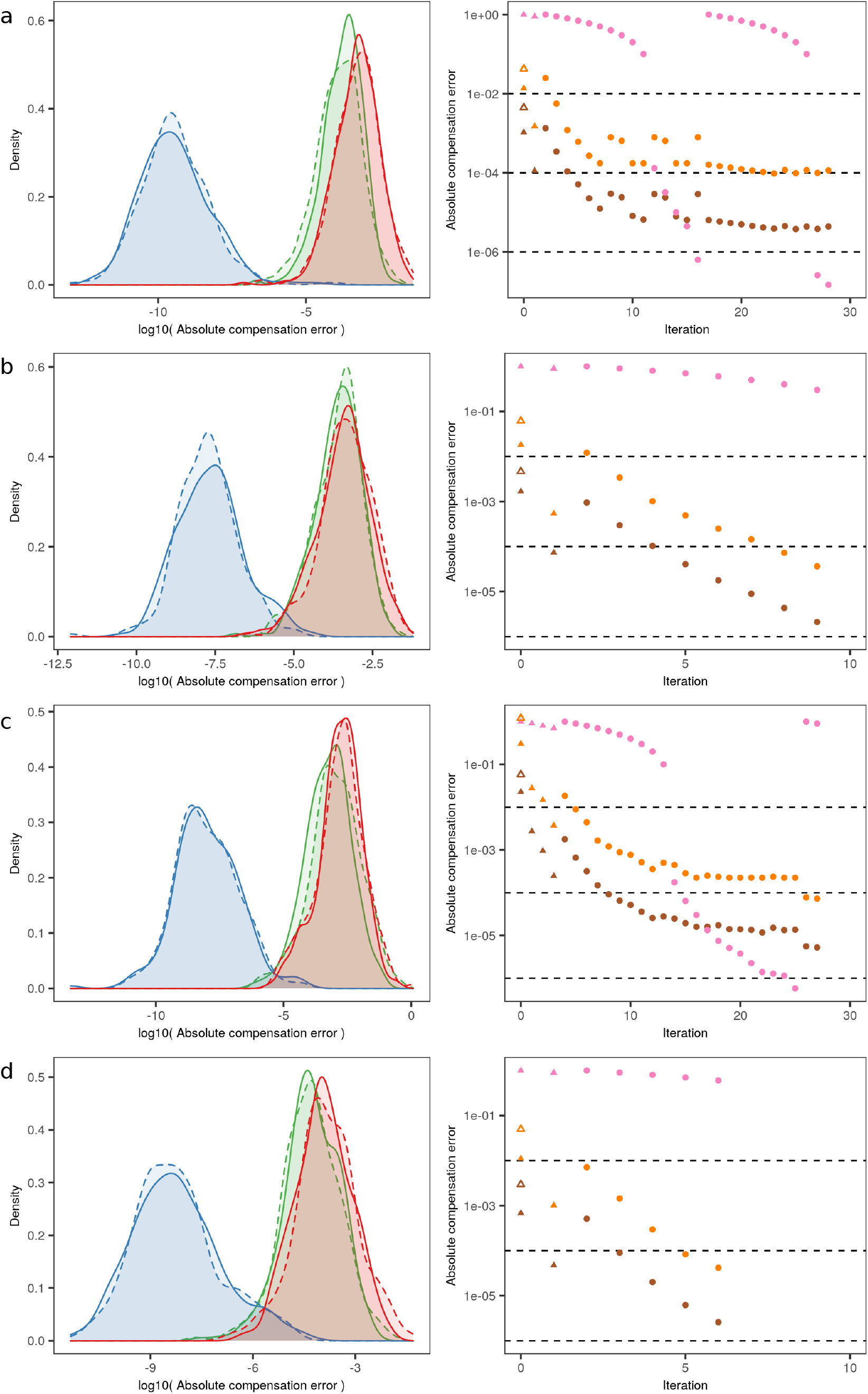
Iterative reduction of compensation error yields optimal spillover coefficients. Each row shows the iterative reduction of compensation error for each dataset: (a) MM1, (b) HS1, (c) HS2, and (d) Be1. Left column displays the densities of compensation error after compensating with positive and negative populations (red), after the first step of AutoSpill (green), and at the final step of AutoSpill (blue). Errors are displayed in log-scale of absolute values, separated in positive (solid lines) and negative (dashed lines) values. Right column displays the convergence of AutoSpill, with points showing standard deviations of errors (brown), maximum absolute error (orange), and the moving average of the decrease in the standard deviation of errors (pink), used to detect oscillations. Linear regressions were carried out in linear scale (triangles) or bi-exponential scale (circles). Empty triangles (same color code) show compensation errors resulting from calculating spillover coefficients with positive and negative populations. Dashed lines display the thresholds for changing from linear to bi-exponential scale (10^−2^, on the maximum absolute error), reaching convergence (10^−4^, on the maximum absolute error), and detecting oscillations (10^−6^, on the moving average of the decrease in the standard deviation of errors).

While effective in most cases, this strategy for reducing compensation error can become compromised when using controls with low fluorescence levels in the primary channel or other fluorescence artifacts. Under these circumstances, iterations gave rise to oscillations in the observed compensation errors before reaching convergence (Fig. 3a,c). In order to deal with these extreme cases, we applied a fraction of the update to the spillover matrix, slowing down convergence and further decreasing compensation error (Fig. 3a,c).

Overall, the iterative refinement of spillover coefficients was effective at reducing errors in compensation. In the four representative datasets reported here, the refinement reduced error from the initial compensation step in 4–6 orders of magnitude (Fig. 3). This low error amounts to optimal spillover coefficients and compensation matrices, relative to the quality of the single-color controls used as input, and therefore it removes a key challenge to successful compensation in high-dimensional flow cytometry.

### Removal of autofluorescence through compensation with an additional autofluorescence channel

Cells produce autofluorescence, due to the interaction of the constituent organic molecules with the incoming photons. The amount of autofluorescence varies between cell types, and it is, for example, higher on cells from the myeloid lineage^13,14^. This can create problems in the analysis of certain flow cytometry datasets. Although the amount of autofluorescence varies between cell types, the spillover from autofluorescence observed in an unstained control (Fig. 4a) behaved similarly to the spillover detected from (exogenous) fluorescent dyes (Fig. 2, first and third column), with the key feature of not having well-defined positive and negative populations. The capacity of AutoSpill to estimate spillover coefficients without needing these populations allowed the treatment of autofluorescence as coming from an endogenous dye, whose single-color control was an unstained control, and whose fluorescence level was recorded in an extra empty channel assigned to a dummy dye. We therefore tested the ability of AutoSpill to compensate out autofluorescence, which was in issue in the HS1 and HS2 datasets. In effect, we were able to use the extra channel to measure the intensity of autofluorescence and greatly reduce its impact onto the other channels (Fig.4b,c). Importantly, the empty channel assigned to autofluorescence worked best when it was the channel with higher level of signal in the unstained control. This way, the most autofluorescent channel was sacrificed during panel design to enhance resolution across all the other channels.

**Figure 4:**
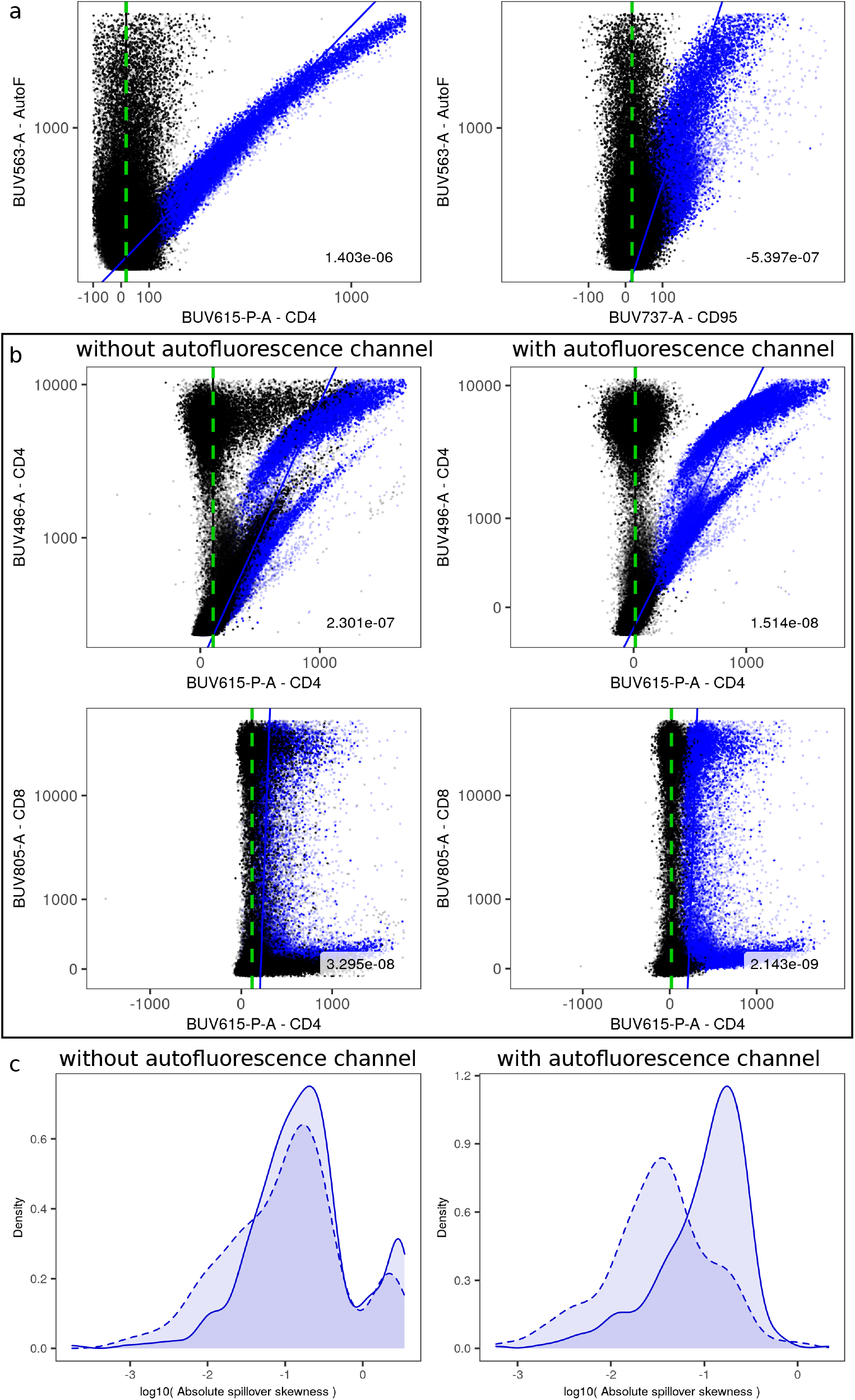
Removal of autofluorescence through compensation with an additional autofluorescence channel. (a) Examples, for the unstained control of the HS1 dataset, of compensation of spillover from the autofluorescence channel (*y*-axes) to two secondary channels (*x*-axes). Uncompensated data is displayed in blue and compensated data in black. Lines show calculated regressions (same color code as data). Resulting compensation errors (slope coefficients of the regressions on compensated data) are shown at the bottom right of each panel. Vertical green dashed lines are shown as a reference for perfectly compensated data. (b) Compensation of two channels (one case per row) severely affected by autofluorescence in the HS1 dataset (left, without autofluorescence channel; right, with autofluorescence channel), with primary channels in *y*-axes and the secondary channels in *x*-axes. Same color and line code, and number with compensation error, as in (a). (c) Density of spillover skewness in the HS1 dataset, without (left) or with (right) autofluorescence channel. Errors are displayed in log-scale of absolute values, separated in positive (solid lines) and negative (dashed lines) values. Autofluorescence, causing spurious positive spillover, corresponds to anomalously large positive skewness in the affected channels (left).

### Linear models for estimation of the Spillover Spreading Matrix

Spillover spreading is defined as the incremental increase in standard deviation of fluorescent intensity in one parameter caused by the increase in fluorescent intensity of another parameter. The SSM coefficients can be calculated by comparing the fluorescent intensity in the primary detector to the standard deviation of fluorescence in the secondary detector, for a pair of positive and negative populations in a single-color control corresponding to the primary detector^15^. It can also be demonstrated the linearity of this relationship for different sizes 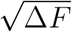, and that the estimation of each spillover spreading coefficient is machine-dependent and compensation-matrix-dependent, but is, however, *dataset independent*^15^. Here, we used quantile partitioning and linear regression to estimate the linear relationship observed by Nguyen *et al*, thereby allowing the inclusion of events above, below, or in-between the positive and negative populations of the original approach.

The events of each single-color control were partitioned quantile-wise in the primary detector, and the standard deviation of the level of autofluorescence was estimated, for each quantile bin, in every secondary detector. Next, two linear regressions were used to estimate, first, the standard deviation at zero fluorescence, and second, the spillover spreading coefficient. Coefficients deemed non-significant using an *F*-test were replaced with zeros, as well as any negative coefficients. The majority of quantiles were, in fact, subsamples of the traditional positive and negative populations, but the inclusion of additional quantiles improved the precision of AutoSpread in estimating spillover spreading effects, because all these events conform to the same linear relationship, assuming that they are on-scale and in the linear range of the flow cytometer (Fig. 5a). As a result, AutoSpread accurately estimated spillover spreading for datasets whose compensation matrices successfully orthogonalized the fluorescent signals present in the single-color controls (Fig. 5b).

**Figure 5:**
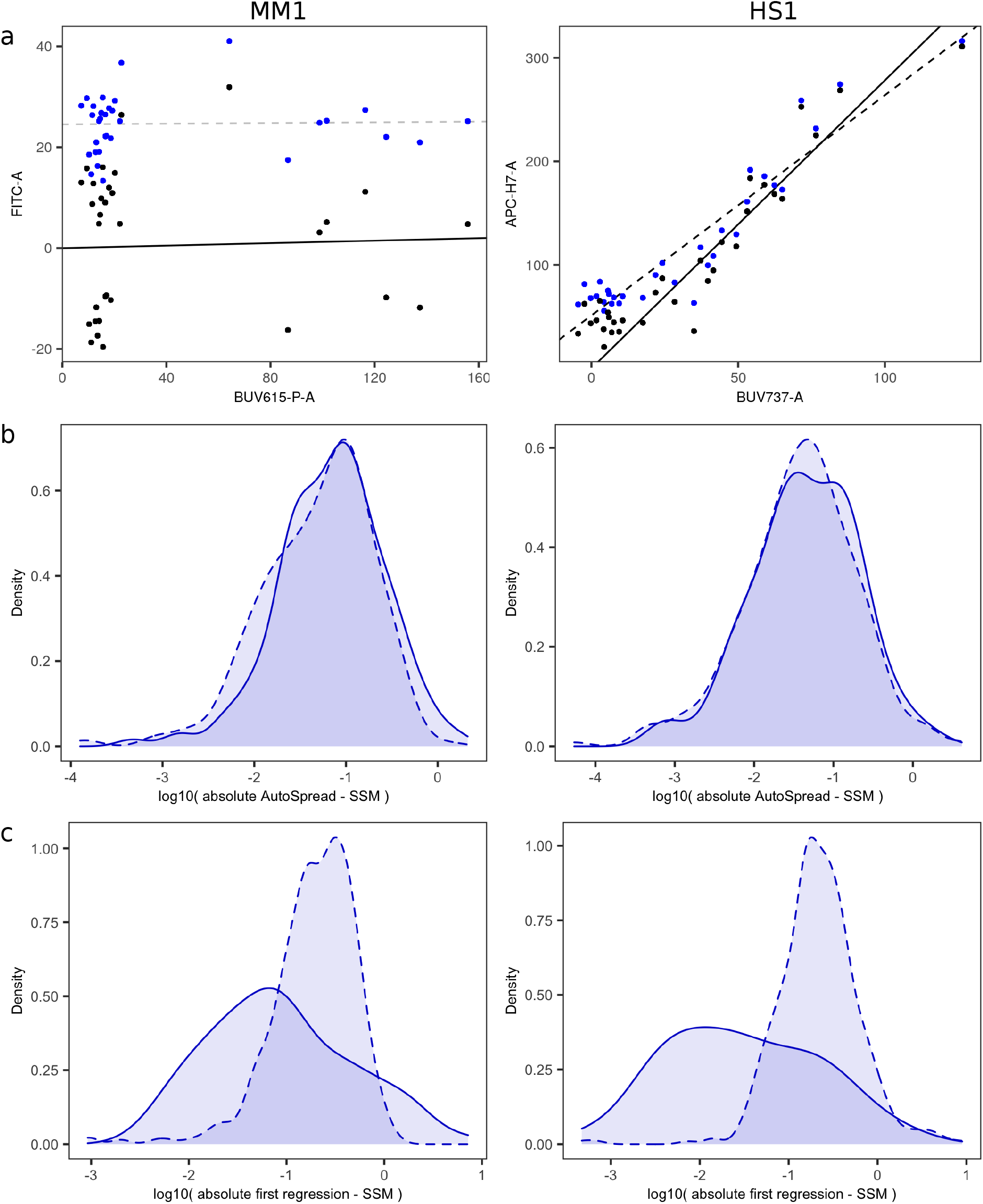
Linear models for estimation of the Spillover Spreading Matrix. Examples are shown for the datasets MM1 (left) and HS1 (right). (a) Regression carried out over the gated events of one single-color control of each dataset, with no well-defined positive and negative populations, with the primary and secondary channels as indicated, respectively, in the *y*- and *x*-axes. Uncompensated data points are displayed in blue and compensated ones in black. Regression from uncompensated (resp. compensated) data is displayed with dashed (resp. solid) lines, in black (resp. gray) when the regression coefficient is significant and positive (resp. non-significant or non-positive). (b) Comparison between the results obtained with AutoSpread vs the usual SSM algorithm, showing the small difference between both calculations. Values are displayed in log-scale of the absolute value of the difference, separated in positive (solid lines) and negative (dashed lines) values. (c) Comparison between results obtained with AutoSpread vs the usual SSM algorithm, but with the omission of the first regression in AutoSpread, which leads to a systematic downward bias in AutoSpread results. Same scale and line code as in (b).

The adjustment step of AutoSpread (the first regression) was critical. The adjustment removed the minor quadratic effect caused by *σ*_0_ in the initial estimates, thereby allowing a more accurate estimation of the coefficients 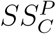. If this adjustment step were skipped, that is, if the *β*’s were taken as the spillover spreading coefficients, then spreading effects would be consistently underestimated. In that case, comparison against the traditional SSM algorithm would show a clear negative bias (Fig. 5c). Including the adjustment step eliminated that bias. For datasets whose single-color controls were contaminated by uncompensated signals (e.g. autofluorescence), both AutoSpread and the traditional SSM calculation may fail to accurately estimate spillover spreading. Initial gating that actively eliminates such effects, as well as the use of an extra autofluorescence channel, can alleviate the problem for both algorithms.

### Biological utility of AutoSpill

To demonstrate the biological utility of improving the spillover matrix, we compared downstream analyses resulting from data compensated with AutoSpill versus the current traditional compensation algorithm. Analyzing 18- and 28-parameter flow cytometry datasets (MM3 and MM2, respectively), we identified multiple examples of poor discrimination of well-described immunological populations due to over- and undercompensation (Fig. 6a). While these errors can readily be identified as compensation errors, AutoSpill also corrected less obvious downstream analyses. For example, in the 18-parameter MM3 dataset, where we gated for CD4+CD8-CD25+ lymphocytes, the population was 10-fold lower using traditional compensation algorithms than with AutoSpill, despite similar compensation identified between the CD4, CD8, and CD25 channels (Fig. 6b). Backgating the missing CD25+ population identified the problem as undercompensation between the CD25 and CD19 channels, leading to elimination of more than 90% of the CD25+ population during early gating stages (Fig. 6c). Finally, we display two clear examples of the benefit of autofluorescence reduction, both based on highly autofluorescent myeloid populations (MM4 & MM5 datasets). First, microglia, a brain-resident macrophage-like population, are often described as having low expression of MHCII during homeostasis^16^. This is a key difference from brain-resident macrophages, with high baseline MHCII expression, and determines the ability of the cell to present antigen to CD4 T cells. Using traditional compensation algorithms, low expression of MHCII was detected on 40% of microglia. This figure, however, dropped to near 0% when autofluorescence reduction was added (Fig. 6d), consistent with the complete absence of MHCII expression at the mRNA level in single cell transcriptome analysis (not shown). We validated the result by including microglia from MHCII knockout mice, where a similar level of background MHCII expression was observed (Fig. 6d), demonstrating that autofluorescence reduction gave the biologically correct outcome. As an independent example, we investigated Foxp3 expression, the key lineage-determining factor of regulatory T cells. Foxp3 expression has also been reported on various autofluorescent lineages, including thymic epithelium^17^, lung epithelium^9^, tumor cells^18^ and macrophages^19^. While expression outside the regulatory T cell lineage was later demonstrated to be due to autofluorescence artifacts ^20-23^, the incorrect reports resulted in research misdirection for several years. Using high dimensional analysis on a Foxp3^GFP^ reporter line and traditional compensation, low expression of the reporter was detected in 10% of the CD11b+ macrophage population (Fig. 6e). This expression was almost entirely eliminated through the use of the autofluorescence correction of AutoSpill, and was validated against wildtype mice, which do not have a GFP reporter present (Fig. 6e). Together, these practical examples demonstrate the added value of AutoSpill to flow cytometry analysis.

**Figure 6:**
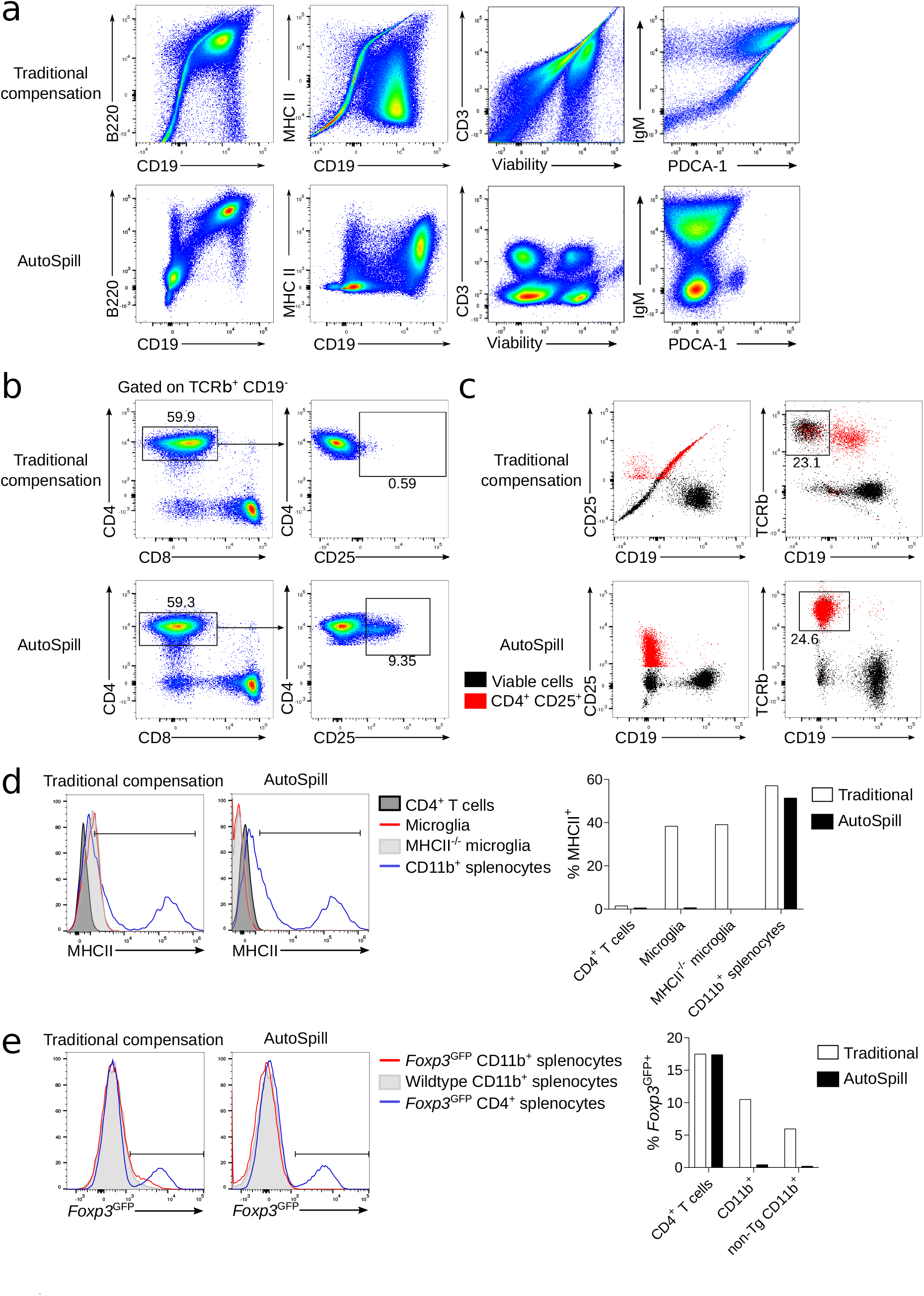
Biological utility of AutoSpill. Downstream analyses of data compensated by either the traditional compensation algorithm or AutoSpill. All plots were prepared from the same FCS files and compensated using FlowJo v.10.7, using either the traditional algorithm or the AutoSpill option. (a) Representative flow cytometry plots illustrating errors corrected by AutoSpill (first and second column, MM3 dataset; third and fourth column, MM2 dataset). (b) Hierarchical gating for CD4+CD8+CD25+ lymphocytes, using data compensated by the traditional algorithm or AutoSpill (MM3 dataset). (c) The CD4+CD25+ population was backgated to identify the source of population loss in the traditional algorithm (MM3 dataset). (d) MHCII expression on known negative cells (CD4 T cells), known positive cells (CD11b+ splenocytes), and microglia (MM4 dataset). Percent positive was thresholded using CD4 T cells as the negative. MHCII knockout microglia were used as a “true negative” staining control. (e) Foxp3^GFP^ expression on known bimodal cells (CD4+ splenocytes) and CD11b+ macrophages (MM5 dataset). The positive population was thresholded using the negative CD4 T cell peak. Wildtype mice, without the GFP transgene, were used as a “true negative” staining control.

## Discussion

Flow cytometry has been a revolutionary force in single-cell analysis. The ability to rapidly analyze protein expression of millions of cells at single-cell level, coupled with the purification capacity of fluorescence-activated cell sorting, has provided a remarkable tool for understanding cellular heterogeneity and function. Initial limitations were overcome through ingenious technical developments: the number of fluorescent parameters were expanded through the development of new dyes and lasers, intracellular staining protocols were optimized for the detection of intracellular (and even post-translationally modified) proteins, RNAflow techniques allowed measurement at the RNA level^24^, and numerous non-antibody-based dyes were able to detect processes from redox potential^25^ to organelle content and status^26^. The very utility of the technique has pushed flow cytometry to its technical barrier —the desire to measure everything on every cell has driven up the number of parameters that can be distinctly measured. The constraints imposed by overlapping fluorescent spectra are arguably the largest limit to the potential of flow cytometry, yet progress in the mathematical underpinnings of the analysis have substantially lagged behind the advances in the chemical and physical bases of the technology.

Newer single-cell technologies, most notably mass cytometry and single-cell RNA-Seq, do not have the spillover issues of flow cytometry. Mass cytometry is a direct competitor to flow cytometry, also primarily utilizing antibody-based detection of single-cell expression^27^. As the heavy metal labels do not overlap, mass cytometry panels can be built up in an modular manner, without the same design constraints required for flow cytometry^28^. While spectral flow cytometry and mass cytometry can readily run more than 40 parameters, classical flow cytometry experiments struggle to use more than 30 parameters, due to the challenge of distinguishing signals from each dye or fluorophore. Nonetheless, flow cytometry has major advantages over mass cytometry, most notably the speed of data acquisition (around 50-fold more rapid data collection) and the ability to sort live cells. The other main competitor to flow cytometry is single-cell RNA-Seq^27^. While initially limited to measurement of RNA content in a semi-quantitative manner, the advent of barcoded antibodies in protocols such as CITE-Seq^29^ and Abseq^30^ provided data directly comparable to that of flow cytometry. As barcoding approaches have no practical limit concerning compensation issues, they can compete with flow cytometry. Even in this case, however, flow cytometry has distinct technological advantages. In addition to the previously mentioned advantage of live-cell sorting, flow cytometry produces data at an unparalleled speed, with more than 10^6^ cells measured per minute, and with a data format enabling immediate analysis. In terms of price, current flow cytometry assays are several orders of magnitude cheaper than RNA-Seq, with costs on the order of 10 USD per 10^6^ cells^27^. Flow cytometry is therefore very much a living technology, with important advantages over competitor technologies and limited only by the parameter barrier.

We have presented here a novel compensation method, which greatly reduces compensation error and expands the possible number of parameters in flow cytometry experiments. The use of robust linear regression and iterative refinement allows the calculation of spillover matrices without the need for using controls with well-defined positive and negative populations, thus permitting the use of the actual panel antibodies for the controls in many experiments. This method can be applied to any flow panel from 4–6 fluorophores up to multi-color staining sets with more than 30 fluorescent dyes. Given that the typical number of gated events in single-color controls is at least in the order of thousands, the amount of data points available enables this approach to reduce compensation errors to such small values that the resulting compensation is, in practical terms, functionally perfect for the given set of single-color controls. On the other hand, the method needs some level of fluorescence in the primary channel for each control (or at least in one of the detectors for spectral systems), to be able to regress the spillover coefficients.

An added feature of AutoSpill is the ability to compensate out autofluorescence. Although some methods have been proposed^31–33^, typically it is not possible to remove autofluorescence, with the exception of some spectral systems^34,35^. By default, AutoSpill does not use an unstained control, but it can be included and assigned to an extra unused channel in the flow cytometer. Data collected in this extra channel can be treated as coming from an endogenous fluorescent dye, which results in the inclusion of autofluorescence levels in the calculation of spillover coefficients and ensuing compensation. This optional approach is recommended when there are non-negligible levels of autofluorescence in one or several channels (as observed from an unstained control), and one of those high-autofluorescence channels is not used in the design of the panel. As autofluorescence can be increased by physiological and cellular processes^13,36^, the ability to compensate out autofluorescence can remove distortions appearing as false positives, where cellular changes are mistakenly identified as altered expression of a marker, while the signal is in fact caused by autofluorescence. This approach will be of particular utility in the study of cell populations with high intrinsic autofluorescence, such as myeloid-lineage cells^13,14^ or tumor cells^37,38^.

AutoSpill focuses on a better estimation of the spillover coefficients, rather than how these coefficients are used to compensate. This distinction is important for spectral flow cytometers^39^, because for those systems different algorithms have been proposed for unmixing ^40,41^, and most of them use single-color controls to estimate the emission spectra of dyes. Each of these spectra is no different from a collection of spillover coefficients, one per channel or detector in the spectral system. In this sense, the method proposed here is equally applicable for a better estimation of the spillover coefficients, that is, of the spectral signatures of dyes in these systems.

In comparison with previous compensation methods, which do not guarantee an upper bound on the compensation error, AutoSpill provides a spillover matrix with such a guarantee, given a set of controls. Therefore, it is possible now to address a new question: To which extent a set of single-color controls is sufficient to ensure proper compensation of data obtained with a complete panel, that is, not just for the set of controls. In our experience, some panels still require minor modifications of the spillover matrix, which implies that the single-color controls do not fully describe the fluorescence properties of the complete panel, probably because of second-order phenomena such as secondary fluorescence or other interactions between dyes. Thus, this remains an open question.

While we demonstrate the utility of this method using eight representative datasets, the tool has been beta-tested more than 1,000 times over a period of 22 months by more than 100 collaborating immunologists. This has allowed the development of a robust algorithm, designed to accommodate diverse datasets and to deal with less-than-perfect data arising in real-world experiments. The code is open source and is released with a permissive license, allowing integration into existing flow cytometry analysis pipelines in academia and industry. To increase access by research communities in immunology and other fields, we also provide a website (https://autospill.vib.be) that allows the upload of sets of single-color controls for calculating the spillover matrix with AutoSpill, produced in formats compatible with common software for flow cytometry analysis. As we have demonstrated by including AutoSpill in FlowJo v.10.7, this algorithm is suitable for integration into commercial software, allowing for rapid and widespread uptake of superior flow cytometry compensation.

## Acknowledgments

This project has received funding from the European Union’s Horizon 2020 research and innovation programme under grant agreement No 779295. This work was also supported by the VIB, the ERC Consolidator Grant TissueTreg (to AL), the Biotechnology and Biological Sciences Research Council through Institute Strategic Program Grant funding BBS/E/B/000C0427 and BBS/E/B/000C0428, and the Biotechnology and Biological Sciences Research Council Core Capability Grant to the Babraham Institute. The authors thank all the collaborators who extensively tested and gave feedback on the beta version of AutoSpill.

## Author Contributions

The study was conceived by CPR and AL. CPR developed and tested the AutoSpill algorithm. OTB, TP, CW, and SHB provided the datasets, input on flow cytometry practicalities, and evaluated compensation results. RH and JS developed and tested the AutoSpread algorithm, and implemented AutoSpill into FlowJo. LK, JC, and AB developed and tested the AutoSpill website. The manuscript was written by CPR and AL, and revised and approved by all authors.

## Competing Financial Interests

The VIB and the Babraham Institute received funding from BD Bioscience in return for pre-publication access to and consultancy on the AutoSpill algorithm, in order to be incorporated into FlowJo v.10.7. RH and JS are affiliated with FlowJo, a wholly owned subsidiary of Becton, Dickinson and Company. The other authors declare no competing financial interests.

## Methods

### Datasets

Collaborating immunologists beta-tested AutoSpill over a period of 22 months, which allowed extensive testing and improvement of the algorithm for niche cases. Among these datasets, four are used as examples here, covering mouse cells, human cells, and beads. Compensation using AutoSpill, with default parameters, was carried out for each of these four sets of single-color controls: mouse splenocytes (MM1 dataset), human PBMCs (HS1 & HS2 datasets), and beads (Be1 dataset). We also analyzed four fully stained datasets, as examples of biological utility: mouse splenocytes (MM2 & MM3 datasets), and mouse microglia (MM4 & MM5 datasets).

### Be1 dataset, beads

UltraComp eBeads™ Compensation Beads (Thermofisher) were used to optimize fluorescence compensation settings for multi-color flow cytometric analysis at a Symphony flow cytometer. UltraComp eBeads™ were stained with the following fluorochrome-labeled anti-human antibodies: anti-CD8–BUV805, anti-CD4–BUV496, anti-CD86–BUV737, anti-CD141–BUV615-P, anti-CD56–BUV563, anti-CD16–BUV395, anti-CD123–BB660-P, anti-CD80–BB630, anti-CD21–BV785, anti-CD27–BV750-P, anti-BAFF-R–BV650, anti-CD94– BV605, anti-CD40–APC-R700 (all BD bioscience); anti-CD3–PerCP-Vio700 (Miltenyi Biotec); anti-CD57–FITC, anti-CD14–PE-Cy5.5, fixable viability dye eFluor780 (all eBioscience); anti-CD24–BV711, anti-CD19–BV480, anti-HLA-DR–BV570, anti-IgM–BV421, anti-CD11c–APC, anti-CD38–PE/Dazzle 594, anti-CD10–PE-Cy5, anti-IgD–PE-Cy7 (all BioLegend).

### HS1 dataset, human PBMCs

Peripheral blood mononuclear cells (PBMC) were isolated from heparinized blood samples of human healthy donors using Ficoll-Paque density centrifugation (MP biomedicals), frozen and then stored in liquid nitrogen. Frozen PBMCs were thawed and counted, and cell concentration was adjusted to 1 × 10^6^ for each single-color control. Cells were plated in a V-bottom 96-well plate, washed once with PBS (Fisher Scientific) and stained with live/dead marker and fluorochrome-conjugated antibodies against surface markers: anti-CD8-BUV805, anti-CD4-BUV496, anti-CD95-BUV737, anti-CD4-BUV615-P, anti-CD28-BB660-P, anti-CD4-BB630, anti-CD4-BV750-P, anti-CD31-BV480, anti-CXCR5–BV650, anti-CD4–PE, anti-CD4-PE-Cy5 (all BD Biosciences); anti-CD3-PerCP-Vio700 (Miltenyi Biotec); anti-CD3-FITC, anti-CD4-PE-Cy5.5, anti-CCR7-PE-Cy7, anti-CD4-eFluor780 (all eBioscience); anti-CD4-BV786, anti-CD4-BV711, anti-CD4-BV605, anti-HLA-DR-BV570, anti-CD127-BV421, anti-CD4-PE/Dazzle 594, anti-CD4-AF647 (all BioLegend).

Samples were stained for 60 min at 4°C, washed twice in PBS/1% FBS (Tico Europe), and then fixed and permeabilized with Foxp3 Transcription Factor Staining Buffer Set (eBioscience), according to manufacturers instructions. Cells were stored overnight at 4°C and were then acquired on a Symphony flow cytometer with Diva software (BD Biosciences). A minimum of 5 × 10^4^ events were acquired for each sample.

### HS2 dataset, human PBMCs

Frozen PBMCs from human healthy donors were processed as for the HS1 datasset and stained with live/dead marker and fluorochrome-conjugated antibodies against the following surface markers: anti-CD8-BUV805, anti-CD4-BUV496, anti-CD95-BUV737, anti-CD28-BB660-P, anti-ICOS-BB630, anti-CXCR3-BV785, anti-PD-1–BV750-P, anti-CXCR5–BV650, anti-CCR2-BV605, (all BD Biosciences); anti-CD3-PerCP-Vio700 (Mil-tenyi Biotec); anti-CD45RA–FITC, anti-CD14–PE-Cy5.5, fixable viability dye eFluor780 (all eBioscience); anti-CD25–BV711, anti-CD31–BV480, anti-HLA-DR–BV570, anti-CD127– BV421, anti-CCR4–PE/Dazzle 594, anti-CCR7–PE-Cy7 (all BioLegend).

Samples were stained for 60 min at 4°C, washed twice in PBS/1% FBS (Tico Europe), and then fixed and permeabilized with Foxp3 Transcription Factor Staining Buffer Set (eBioscience), according to manufacturers instructions. Cells were stained overnight at 4°C with anti-Ki67–BUV615-P, anti-CTLA-4–PE-Cy5, anti-RORγt–PE (BD Biosciences) and anti-FOXP3–AF647 (BioLegend) anti-human intracellular antibody. Samples were acquired on a Symphony flow cytometer (BD Biosciences).

### MM1 dataset, mouse splenocytes

Splenocytes from C57Bl/6 mice were disrupted with glass slides, filtered through 100 *μ*m mesh, and red blood cells lysed. Cells were fixed and permeabilized with Foxp3 Transcription Factor Staining Buffer Set (eBioscience) according to the manufacturer’s instructions, and stained overnight at 4°C with Fixable Viability Dye eFluor780 (eBioscience) or the following antibodies: anti-CD4–BV421, anti-CD24–BV510, anti-CD3–BV570, anti-CD4–BV605, anti-CD3–BV650, anti-CD4–BV711, anti-CD4–BV785, anti-CD3–AF488/anti-CD4–AF488/anti-TCR*β*–AF488, anti-CD4–PerCP-Cy5.5, anti-CD4–PE-594, anti-CD8–PE-Cy7, anti-MHC-II–AF700 (all Biolegend), anti-CD19–BV750, anti-CD3–BB630-P/anti-Thy1.2–BB630-P, anti-CD45.2–BB660-P2/anti-CD3–BB660-P2, anti-TCRβ–BB790-P, anti-CD4–BUV395, anti-IgD–BUV496, anti-CD3–BUV563, anti-CD3–BUV615-P, anti-CD19–BUV661, anti-CD21–BUV737, anti-CD8–BUV805 (all BD Biosciences), anti-CD4–PE/anti-CD3–PE/anti-CD8–PE, anti-IgM–PE-Cy5, anti-CD3–PE-Cy5.5 or anti-CD4–APC (all eBioscience). For some fluorophores, multiple antibodies were used in the same compensation control, which is indicated by slashes. Samples were acquired on a Symphony flow cytometer (BD Biosciences).

### MM2 dataset, mouse splenocytes

Splenocytes from C57Bl/6 mice were disrupted with glass slides, filtered through 100 *μ*m mesh, and red blood cells lysed. Cells were stained with Fixable Viability Dye eFluor780 (eBioscience), fixed and permeabilized with Foxp3 Transcription Factor Staining Buffer Set (eBioscience) according to the manufacturer’s instructions, and stained overnight at 4°C with the following antibodies: anti-CD4–BV421, anti-CD24–BV510, anti-Ly6G–BV570, anti-XCR1–BV650, anti-CD19–BV785, anti-CD3–AF488, anti-PDCA-1–PerCP-Cy5.5, anti-CD23-PE, anti-CD64–PE-594, anti-CD172a–PE-Cy7, anti-CD45–APC, anti-MHCII–AF700 (all Biolegend), anti-IgE–BV605, anti-CD93–BV711, anti-CD11b–BV750, anti-CD80–BB630-P, anti-CD95–BB660-P2, anti-TCR*β*–BB790-P, anti-CD103–BUV395, anti-IgD–BUV496, anti-Ly6C–BUV563, anti-Siglec F–BUV615-P, anti-c-Kit–BUV661, anti-CD21/35–BUV737, anti-CD8a–BUV805 (all BD Biosciences), anti-IgM–PE-Cy5 and anti-NK1.1–PECy5.5 (eBioscience). Compensation controls were stained as described in the MM1 dataset. Samples were acquired on a Symphony flow cytometer (BD Biosciences).

### MM3 dataset, mouse splenocytes

Splenocytes from C57Bl/6 mice were disrupted with glass slides, filtered through 100 *μ*m mesh, and red blood cells lysed. Cells were stained with Fixable Viability Dye eFluor780 (eBioscience), anti-CD90.2–BV510, anti-CD25–BV650, anti-CD45–BUV395 (all Biolegend), anti-CD127–PE and anti-B220–PE-Cy5 (all eBioscience). Cells were fixed and permeabilized with Foxp3 Transcription Factor Staining Buffer Set (eBioscience) according to the manufacturer’s instructions, and stained overnight at 4°C with the following antibodies: anti-T-bet–BV421, anti-CD8–BV785, anti-NKp46–FITC, anti-NK1.1–PE-Cy5.5, anti-MHCII-AF700 (all Biolegend), anti-CD11b–eFluor450, anti-GATA3–PE-Cy7, anti-CD3–biotin, anti-RORt–APC (all eBioscience), anti-TCR*β*–BB790-P, anti-CD4–BUV496 and anti-CD19-BUV661 (all BD Biosciences). Antibodies used for compensation controls were anti-CD25-BV421, anti-CD44-BV510, anti-CD3-BV650, anti-CD8-BV785, anti-NK1.1–PE-Cy5.5, anti-MHCII–AF700 (all Biolegend), anti-CD11b–eFluor450, anti-TCR*β*–FITC, anti-B220–PE-Cy5, anti-CD23–PE-Cy7, anti-CD8–biotin, anti-Foxp3–APC, anti-CD69–PE (all eBioscience), anti-TCRβ–BB790-P, anti-CD103–BUV395, anti-CD4–BUV496 and anti-CD19–BUV661 (all BD Biosciences). Streptavidin AF350 (Invitrogen) was used to identify biotinylated antibody. Samples were acquired on a Yeti/ZE5 flow cytometer (Propel Labs/BioRad).

### MM4 dataset, mouse microglia

MHCII knockout mice^42^ were used on the B6 background. Leukocytes and microglia were extracted from mouse brains by chopping with a razor blade, digest in 0.4mg/ml collagenase D (Sigma-Aldrich) and separation over 40% Percoll (GE Healthcare). Microglia were stained with anti-MHCII–FITC (clone M5/114.15.2, eBioscience), anti-CD11b–PE-Cy7 (clone M1/70, eBioscience), anti-CD45–APC (clone 30-F11, eBioscience), anti-CD4–PE-Dazzle594 (clone GK1.5, BioLegend) and fixable viability dye eFluor780 (eBioscience).

### MM5 dataset, mouse microglia

Foxp3^DTR–GFP^ mice^43^ were used on the B6 background. Leukocytes and microglia were extracted from mouse brains by chopping with a razor blade, digest in 0.4mg/ml collagenase D (Sigma-Aldrich) and separation over 40% Percoll (GE Healthcare). Microglia were stained with anti-MHCII–FITC (clone M5/114.15.2, eBioscience), anti-CD11b–PE-Cy7 (clone M1/70, eBioscience), anti-CD45–APC (clone 30-F11, eBioscience), anti-CD4–PE-Dazzle594 (clone GK1.5, BioLegend) and fixable viability dye eFluor780 (eBioscience).

### General implementation details of AutoSpill

AutoSpill was implemented in R v.3.6.3, using the packages flowCore v.1.52.1, flowWorkspace v.3.34.1, ggplot2 v.3.3.2, moments v.0.14, and RColorBrewer v.1.1-2. Further details on packages specific to particular steps of the algorithm are listed below.

### Initial gating

The initial gate was calculated independently for each control, over the 2d-density of events on forward and side scatter (FSC-A and SSC-A parameters). To robustly detect the population of interest, two tessellations were successively carried out to isolate the desired density peak. First, data was trimmed on extreme values (1% and 99%). Then, maxima were located numerically by a moving average (window size 3) on a soft estimation of the 2d-density (bandwidth factor 3). The first tessellation was carried out on these density maxima, and the tile corresponding to the highest maximum was selected, ignoring peaks with lower values of both FSC-A and SSC-A (less than 5% of range). A rectangular region in the FSC-A/SSC-A-plane was chosen by using the median and 3× the mean absolute deviation of the events contained in the selected tile. A second, finer 2d-density estimation (bandwidth factor 2) was obtained on this region, followed again by numerical detection of maxima (window size 2) and tessellation by the maxima. A final 2d-density estimation (bandwidth factor 1) was obtained on the tile containing the highest maximum, with the gate being defined as the convex hull enclosing the points that belonged to this tile and had a density larger than a threshold (33% of range).

Tessellations were carried out with package deldir v.0.1-28, density estimations with packages MASS v.7.3-51.6, surface interpolations with package fields v.10.3, and spatial operations with packages sp v.1.4-2 and tripack v.1.3-9.

### Robust linear models for estimation of spillover coefficients

The linearity of the quantum mechanical nature of photons implies that the ratio between the average fluorescence level (that is, the average number of photons) detected in any two detectors and from any dye is equal to the ratio between the corresponding values of the emission spectrum of the dye, regardless of the level of fluorescence. As the value of the spillover coefficient for the primary channel (the channel assigned to the dye in the single-color control, in classical systems) is usually normalized to one, the spillover coefficient of every secondary channel is equal to the fluorescence ratio above. This implies that each spillover coefficient can be directly read from the slope of a linear regression considering the fluorescence in the primary channel as the independent variable and the fluorescence in the secondary channel as the dependent variable (that is, with *x* and *y* swapped for the usual representation of single-color controls when compensating). Thus, absence of spillover corresponds to a zero slope in this regression, that is, to the vertical direction in the usual plot were the primary channel is displayed in the *y*-axis. To protect the algorithm against distortions in the data, specially those coming from autofluorescence issues, robust linear regression was used, giving lower weights to events farther away from the estimated regression line. Robust linear models were implemented with the package MASS v.7.3-51.6, with default parameters.

### Refinement of spillover matrix

After the first iteration of the algorithm, applying on compensated data the same kind of calculation used for the spillover coefficients, on channels in classical systems or on dyes in spectral systems, would produce zero values with perfect compensation, corresponding to perfectly vertical compensation plots. Otherwise, errors in compensation would yield non-zero values reflecting residual spillover. Overcompensated data would amount to excessively negative values in the secondary channel/dye, corresponding to a negative slope. Similarly, undercompensation would produce excessively positive values in the secondary channel/dye, corresponding to a positive slope.

Observed errors in compensation arise from errors in the estimation of the spillover coefficients. Crucially, it can be proved that, for the average event at any level of fluorescence, the error matrix **T** in the calculation of the spillover matrix **S** can be calculated from the observed compensation errors **E** as:

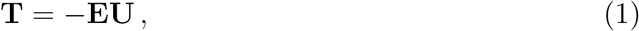

**U** = **S** + **T** being the (erroneous) spillover matrix used to compensate the data (see below).

By successively applying equation Eq. (1), that is, by iteratively refining the spillover matrix and recalculating the compensation, errors in the spillover matrix and errors in compensation can be reduced to a negligible magnitude. The algorithm starts working in linear scale, and switches to bi-exponential scale when the maximum compensation error across all single-color controls is less than a threshold fixed a priori (10^−2^). To be used in Eq. (1), compensation errors obtained in bi-exponential scale are transformed back to linear scale, by using the two points in the regression line with extreme values in the primary channel. Iterations stop near convergence of the algorithm, when the maximum compensation error across all single-color controls is less than a threshold of 10^−4^.

While effective in most cases, this strategy for reducing compensation error can become compromised when using controls with low fluorescence levels in the primary channel or other fluorescence artifacts. In these situations, iterations can give rise to oscillations in the observed compensation errors before reaching convergence. To deal with these extreme cases, oscillations are detected by a moving average (size 10, initial value 1) of the decrease in the standard deviation of spillover errors. When this moving average gets below a threshold of 10^−6^, a fraction (10%) of the update to the spillover matrix is applied in Eq. (1), slowing down convergence and further decreasing compensation error.

### Spillover error

In a flow cytometry system with *c* channels, let us consider the spillover matrix for a set of *d* single-color controls, that is for *d* dyes, with *d* ≤ *c*. We concentrate on the dye *i* = 1… *d* during the following argument.

For any event in the flow cytometer, we have the following two row vectors: the *true* event data **x**, with length *d*, and the *observed* event data **y**, with length *c*. On average for any level of fluorescence, true and observed events are related linearly through the *d* × *c* spillover matrix **S**, according to

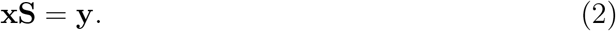

Classical flow cytometry systems have *c = d*, and compensation is usually achieved by inverting the spillover matrix **S** and multiplying by the observed data **y**. Spectral systems feature *c > d*, and compensation is usually called unmixing and is not unequivocally defined, because Eq. (2) produces an overspecified system of equations. In the following, and for simplicity, we refer to unmixing in spectral systems also as compensation.

Independently of the compensation method used, when the spillover matrix **S** is estimated as **U** = **S**+**T**, thus with some error **T**, it unavoidably gives rise to incorrectly compensated data **x** + **p**, which verifies, on average,

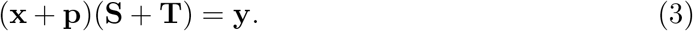

Therefore,

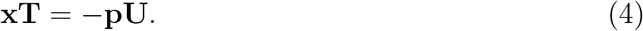

The vectors **x** and **p**, and the matrices **S, T**, and **U**, have the following properties:

- Because **x** represents the true value of events in the single-color control for dye *i*, then *x_i_* > 0 and *X_j_* = 0, for all *j ≠ i*.
- The *i*-th row of the spillover matrix **S** is normalized with 1 = *S_ir_* ≥ *S_is_* ≥ 0, for some *r* = 1… *c* and every *s* ≠ *r*.
- The row normalization of **S** implies that the true value of the dye in the control, *X_i_*, can always be obtained from the observed value *y_r_*, as Eq. (2) implies *y_r_ = x_i_S_ir_ = x_i_*. Therefore, *p_i_* = 0, irrespective of errors in the estimation of the spillover matrix.
- Also because of the row normalization of the spillover matrix, the estimation of the spillover coefficient *S_ir_* = 1 will always be exact, i.e. *U_ir_* = 1 and *T_ir_* = 0, irrespective of errors in the estimation of the spillover matrix.

Let us consider now the LHS of Eq. (4), i.e. the row vector **xT**. Its *s*-th coefficient, for any *s* = 1… *c*, equals

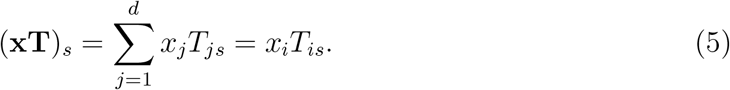

Note that (**xT**)_*r*_ = 0.

Let us consider the RHS of Eq. (4), i.e. the row vector −**pU**. Its *s*-th coefficient, for any *s* = 1… *c*, equals

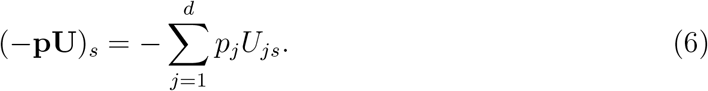

Note that the summation term *p_i_U_is_* = 0.

Equations (4)–(6) imply that, for any *s* = 1… *c*,

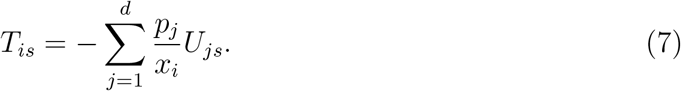

The ratio *p_j_/x_i_* can be considered as the compensation error for the average event, corresponding to a spurious signal assigned to dye *j*, caused by incorrectly compensated spillover from dye *i*. Equation (3) implies that the ratio *p_j_/x_i_* is invariant w.r.t. the level of fluorescence, and thus it can be estimated by regressing *p_j_* vs *x_i_*.

Let us define the compensation error matrix **E** as the *d* × *d* matrix with coefficients

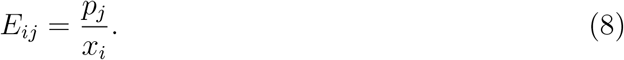

Note that *E_ii_* = 0. We can then rewrite Eq. (7) as

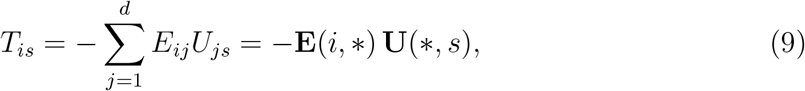

for any *s* = 1… *c*.

In summary, Eq. (9) allows to calculate the *i*-th row of the spillover error matrix **T**. By repeating the same argument for every dye, we can obtain all the rows *i* = 1… *d*, and thus the complete matrix as

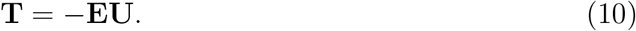

### Refinement of the spillover matrix

The algorithm calculates a first approximation to the spillover matrix, and then it refines it iteratively by successively applying Eq. (10). As before, we refer to unmixing in spectral systems as compensation.

Input: Collection of *d* single-color controls (one per dye) {**Y**_*i*_}, *i* =1… *d*, each one being a matrix with *n_i_* rows (events) and *c* columns (channels), *c ≥ d*.

Output: Spillover matrix **S**, a matrix with *d* rows (dyes) and *c* columns (channels), and the collection of compensated controls {**X**_*i*_}, *i* = 1… *d*, each one being a matrix with *n_i_* rows (events) and d columns (dyes).

Parameters: Upper bound *ϵ* in the compensation error required to achieve convergence.

Algorithm:

1. For each single-color control **Y**_*i*_, *i* = 1… *d*,

For each channel *j* = 1… *c, j ≠ h_i_, h_i_* being the channel with highest signal for dye *i*,

Calculate robust linear model **Y**_*i*_(∗, *j*) ~ **Y**_*i*_(∗, *h_i_*) and obtain slope 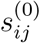.
2. Build initial spillover matrix **S**^(0)^ as

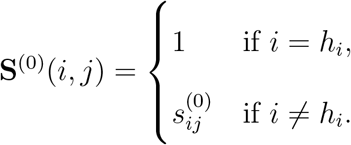
3. Obtain initially compensated controls 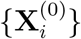, by applying the algorithm of choice with **S**^(0)^ on the controls {**Y**_*i*_}.
4. Refine spillover matrix **S**^(*t*)^ and compensated controls 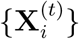, obtaining spillover matrix **S**^(*t*+1)^ and compensated controls 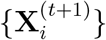, until convergence.

4.1. For each compensated single-color control 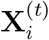, *i* = 1… *d*,

For each other dye *j* = 1… *d, j = i*,

Calculate robust linear model 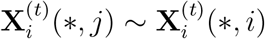 and obtain slope 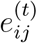.
4.2. Build matrix of compensation errors **E**^(*t*)^ as

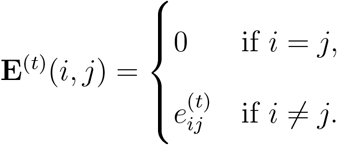
4.3. Calculate non-normalized spillover matrix 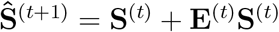.
4.4. Calculate normalized spillover matrix **S**^(*t*+1)^ by rows, as

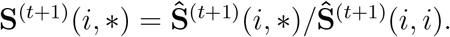
4.5. Obtain compensated controls 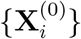, by applying the algorithm of choice with **S**^(*t*+1)^ on the initial controls {**Y**_*i*_}.
4.6. Convergence is attained when ‖**E**^(*t*)^‖ < *ϵ*.
5. At convergence, *t*^∗^ being the last iteration, obtain final spillover matrix and compensated controls as

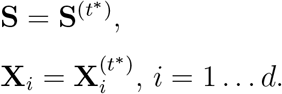

### Linear models for estimation of SSM

Successful compensation equilibrates around zero the fluorescence levels in all secondary channels, but with the cost of introducing undesirable variance or spread in those channels. Again for quantum mechanical reasons, the spread in fluorescence for any (compensated or uncompensated) channel/dye grows linearly with the fluorescence level, and therefore the coefficients of the SSM can be estimated with linear regression.

We start with the formula for an SSM coefficient 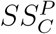, which characterizes the incremental standard deviation induced in parameter *C* by the spillover from parameter *P*^15^,

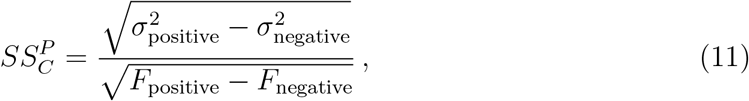

where *σ*_positive_ and *σ*_negative_ are the standard deviations in C-fluorescence in a positive and negative population, respectively, and *F*_positive_ − *F*_negative_ is the difference in P-fluorescence intensity between them. While the traditional algorithm estimates the above quantities using medians and robust standard deviations of fluorescence in the positive and negative populations, we will, for the sake of linear regression, let our negative be the theoretical quantity when *P*-fluorescence (*F*) is equal to zero, while the standard deviation is an unknown quantity, which we call *σ*_0_. This gives us the following equation relating *F* to *σ*, which is suitable for estimating *σ*_0_ by linear regression:

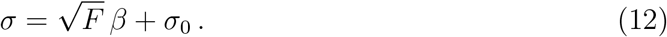

Notice that the slope *β* is not equal to the spillover spreading coefficient 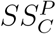, except in the unique case where *σ*_0_ equals zero. We thus proceed with the estimation of *σ*_0_ as the first step of AutoSpread.

To supply data for the regression, we partition the events of the single-color control for parameter *P* by quantile. For controls with a large number of events, we use 256 quantiles, but we allow as few as 8 to ensure enough events in each quantile to estimate standard deviation reliably. For each other parameter *C*, we calculate in each quantile the robust standard deviation of fluorescence (the 84^th^ percentile minus the median) as the estimate of σ, and the median fluorescence as the estimate of *F*. The *F* values may be negative and/or close to zero, so they are passed through a square-root-like transform defined by 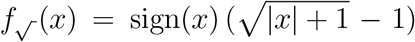 prior to regression, instead of the simple square root function. The resulting regression provides an estimate of *σ*_0_.

Using the estimate of *σ*_0_, AutoSpread calculates for each quantile the estimate of *σ*′, defined by 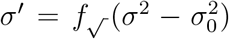, and these adjusted standard deviation estimates provide the data for the second regression, 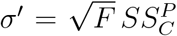. This regression is calculated without an intercept term because the adjustment of *σ***0** forces it to zero.

### Data and Code Availability

The raw data for the eight analyzed datasets is available at FlowRepository (https://flowrepository.org), with ids FR-FCM-Z2SV (Be1), FR-FCM-Z2ST (HS1 & HS2), FR-FCM-Z2SS (MM1), FR-FCM-Z2SW (MM2), FFR-FCM-Z2SJ (MM3), FR-FCM-Z2SK (MM4), and FR-FCM-Z2SL (MM5). Note that the compensation controls for the MM2 dataset are the MM1 dataset.

Source code for AutoSpill is available through the R package autospill, available at the github repository https://github.com/carlosproca/autospill, which includes batch code that reproduces the reported results for the datasets MM1, HS1, HS2, and Be1. The R package is also available in the Supplementary Information as Supplementary Data. In addition, AutoSpill is accessible as a freely-available web service at https://autospill.vib.be. The R package also includes batch code to reproduce results as generated by the website.

To allow a large user base to take immediate advantage of the new approaches reported here, an implementation of AutoSpill is included in the release of FlowJo v.10.7. AutoSpread is available in binary form in FlowJo v.10.7 (patent pending).

## Reference

[1] Herzenberg, L. A. et al. The history and future of the Fluorescence Activated Cell Sorter and flow cytometry: A view from Stanford. Clinical Chemistry 48, 1819–1827 (2002).

[2] O’Gorman, M. R. Clinically relevant functional flow cytometry assays. Clinics in laboratory medicine 21, 779–94 (2001).

[3] Krutzik, P. O. & Nolan, G. P. Fluorescent cell barcoding in flow cytometry allows high-throughput drug screening and signaling profiling. Nature Methods 3, 361–368 (2006).

[4] Maciorowski, Z., Chattopadhyay, P. K. & Jain, P. Basic Multicolor Flow Cytometry. Current Protocols in Immunology 117, 5.4.1–5.4.38 (2017).

[5] Cossarizza, A. et al. Guidelines for the use of flow cytometry and cell sorting in immunological studies (second edition). European Journal of Immunology 49, 1457–1973 (2019).

[6] Bendall, S. C., Nolan, G. P., Roederer, M. & Chattopadhyay, P. K. A deep profiler’s guide to cytometry. Trends in Immunology 33, 323–332 (2012).

[7] Roederer, M. Spectral compensation for flow cytometry: Visualization artifacts, limitations, and caveats. Cytometry 45, 194–205 (2001).

[8] Carr, E. J. et al. The cellular composition of the human immune system is shaped by age and cohabitation. Nature Immunology 17, 461–468 (2016).

[9] Mair, F. & Prlic, M. OMIP044: 28color immunophenotyping of the human dendritic cell compartment. Cytometry Part A 93, 402–405 (2018).

[10] Brummelman, J. et al. Development, application and computational analysis of high-dimensional fluorescent antibody panels for single-cell flow cytometry. Nature Protocols 14, 1946–1969 (2019).

[11] Bandura, D. R. et al. Mass cytometry: Technique for real time single cell multitarget immunoassay based on inductively coupled plasma time-of-flight mass spectrometry. Analytical Chemistry 81, 6813–6822 (2009).

[12] Bagwell, C. B. & Adams, E. G. Fluorescence spectral overlap compensation for any number of flow cytometry parameters. Annals of the New York Academy of Sciences 677, 167–84 (1993).

[13] Mitchell, A. J. et al. Technical Advance: Autofluorescence as a tool for myeloid cell analysis. Journal of Leukocyte Biology 88, 597–603 (2010).

[14] Vermaelen, K. & Pauwels, R. Accurate and simple discrimination of mouse pulmonary dendritic cell and macrophage populations by flow cytometry: Methodology and new insights. Cytometry Part A 61, 170–177 (2004).

[15] Nguyen, R., Perfetto, S., Mahnke, Y. D., Chattopadhyay, P. & Roederer, M. Quantifying spillover spreading for comparing instrument performance and aiding in multicolor panel design. Cytometry Part A 83A, 306–315 (2013).

[16] Li, Q. & Barres, B. A. Microglia and macrophages in brain homeostasis and disease. Nature Reviews Immunolology 18, 225–242 (2018).

[17] Chang, X. et al. The Scurfy mutation of FoxP3 in the thymus stroma leads to defective thymopoiesis. Journal of Experimental Medicine 202, 1141–1151 (2005).

[18] Zuo, T. et al. FOXP3 Is an X-Linked Breast Cancer Suppressor Gene and an Important Repressor of the HER-2/ErbB2 Oncogene. Cell 129, 1275–1286 (2007).

[19] Manrique, S. Z. et al. Foxp3-positive macrophages display immunosuppressive properties and promote tumor growth. Journal of Experimental Medicine 208, 1485–1499 (2011).

[20] Liston, A. et al. Lack of Foxp3 function and expression in the thymic epithelium. Journal of Experimental Medicine 204, 475–480 (2007).

[21] Li, F. et al. Autofluorescence contributes to false-positive intracellular Foxp3 staining in macrophages: A lesson learned from flow cytometry. Journal of Immunological Methods 386, 101–107 (2012).

[22] Kim, J. et al. Cutting Edge: Depletion of Foxp3 + Cells Leads to Induction of Autoimmunity by Specific Ablation of Regulatory T Cells in Genetically Targeted Mice. The Journal of Immunology 183, 7631–7634 (2009).

[23] Put, S. et al. Macrophages have no lineage history of Foxp3 expression. Blood 119, 1316–1318 (2012).

[24] Hanley, M. B., Lomas, W., Mittar, D., Maino, V. & Park, E. Detection of Low Abundance RNA Molecules in Individual Cells by Flow Cytometry. PLoS ONE 8 (2013).

[25] Li, R., Jen, N., Yu, F. & Hsiai, T. K. Assessing mitochondrial redox status by flow cytometric methods: Vascular response to fluid shear stress. Current Protocols in Cytometry 58, 9.37.1–9.37.14 (2011).

[26] Poot, M., Gibson, L. L. & Singer, V. L. Detection of apoptosis in live cells by Mito-Tracken(TM) red CMXRos and SYTO dye flow cytometry. Cytometry 27, 358–364 (1997).

[27] Chattopadhyay, P. K., Winters, A. F., Lomas, W. E., Laino, A. S. & Woods, D. M. High-Parameter Single-Cell Analysis. Annual Review of Analytical Chemistry 12, 411–430 (2019).

[28] Spitzer, M. H. & Nolan, G. P. Mass Cytometry: Single Cells, Many Features. Cell 165, 780–791 (2016).

[29] Stoeckius, M. et al. Simultaneous epitope and transcriptome measurement in single cells. Nature Methods 14, 865–868 (2017).

[30] Shahi, P., Kim, S. C., Haliburton, J. R., Gartner, Z. J. & Abate, A. R. Abseq: Ultrahigh-throughput single cell protein profiling with droplet microfluidic barcoding. Scientific Reports 7 (2017).

[31] Roederer, M. & Murphy, R. F. Cellbycell autofluorescence correction for low signaltonoise systems: Application to epidermal growth factor endocytosis by 3T3 fibroblasts. Cytometry 7, 558–565 (1986).

[32] Alberti, S., Parks, D. R. & Herzenberg, L. A. A single laser method for subtraction of cell autofluorescence in flow cytometry. Cytometry 8, 114–119 (1987).

[33] Roederer, M. Distributions of autofluorescence after compensation: Be panglossian, fret not. Cytometry Part A 89, 398–402 (2016).

[34] Nitta, N., Veltri, G. & Dessing, M. Method and theory of the autofluorescence unmixing in sp6800 spectral cell analyzer. Tech. Rep., Sony Corporation (2015).

[35] Schmutz, S., Valente, M., Cumano, A. & Novault, S. Spectral cytometry has unique properties allowing multicolor analysis of cell suspensions isolated from solid tissues. PLoS ONE 11 (2016).

[36] Surre, J. et al. Strong increase in the autofluorescence of cells signals struggle for survival. Scientific Reports 8 (2018).

[37] Smith, C. A., Pollice, A., Emlet, D. & Shackney, S. E. A simple correction for cell autofluorescence for multiparameter cell-based analysis of human solid tumors. Cytometry Part B - Clinical Cytometry 70, 91–103 (2006).

[38] Pantanelli, S. M. et al. Differentiation of malignant B-lymphoma cells from normal and activated T-cell populations by their intrinsic autofluorescence. Cancer Research 69, 4911–4917 (2009).

[39] Nolan, J. P. & Condello, D. Spectral flow cytometry. Current Protocols in Cytometry 63, 1.27.1–1.27.13 (2013).

[40] Novo, D., Grégori, G. & Rajwa, B. Generalized unmixing model for multispectral flow cytometry utilizing nonsquare compensation matrices. Cytometry Part A 83 A, 508–520 (2013).

[41] Futamura, K. et al. Novel full-spectral flow cytometry with multiple spectrally-adjacent fluorescent proteins and fluorochromes and visualization of in vivo cellular movement. Cytometry Part A 87, 830–842 (2015).

[42] Madsen, L. et al. Mice lacking all conventional MHC class II genes. Proceedings of the National Academy of Sciences of the United States of America 96, 10338–10343 (1999).

[43] Kim, J. M., Rasmussen, J. P. & Rudensky, A. Y. Regulatory T cells prevent catastrophic autoimmunity throughout the lifespan of mice. Nature Immunology 8, 191–197 (2007).

